# Novel regulators of PrP^C^ biosynthesis revealed by genome-wide RNA interference

**DOI:** 10.1101/2021.01.28.428672

**Authors:** Daniel Heinzer, Merve Avar, Daniel Patrick Pease, Ashutosh Dhingra, Jiang-An Yin, Elke Schaper, Berre Doğançay, Marc Emmenegger, Anna Spinelli, Kevin Maggi, Andra Chincisan, Simon Mead, Simone Hornemann, Peter Heutink, Adriano Aguzzi

## Abstract

The cellular prion protein PrP^C^ is necessary for prion replication, and its reduction greatly increases life expectancy in animal models of prion infection. Hence the factors controlling the levels of PrP^C^ may represent therapeutic targets against human prion diseases. Here we performed an arrayed whole-transcriptome RNA interference screen to identify modulators of PrP^C^ expression. We cultured human U251-MG glioblastoma cells in the presence of 64’752 unique siRNAs targeting 21’584 annotated human genes, and measured PrP^C^ using a one-pot fluorescence-resonance energy transfer immunoassay in 51’128 individual microplate wells. This screen yielded 743 candidate regulators of PrP^C^. When downregulated, 563 of these candidates reduced and 180 enhanced PrP^C^ expression. Recursive candidate attrition through multiple secondary screens yielded 54 novel regulators of PrP^C^, 9 of which were confirmed by CRISPR interference as robust regulators of PrP^C^ biosynthesis and degradation. The phenotypes of 6 of the 9 candidates were inverted in response to transcriptional activation using CRISPRa. The RNA-binding post-transcriptional repressor Pumilio-1 was identified as a potent limiter of PrP^C^ expression through the degradation of *PRNP* mRNA. Because of its hypothesis-free design, this comprehensive genetic-perturbation screen delivers an unbiased landscape of the genes regulating PrP^C^ levels in cells, most of which were unanticipated, and some of which may be amenable to pharmacological targeting in the context of antiprion therapies.

## Introduction

A feature common to all prion diseases is the conversion of the cellular prion protein (PrP^C^) into a misfolded, disease-causing isoform called PrP^Sc^ (Aguzzi and De Cecco, 2020). PrP^C^ is an agonist of the adhesion G protein-coupled receptor Adgrg6 in the peripheral nervous system, but its role in the central nervous system (CNS) has remained unclear (Küffer et al., 2016; Wulf et al., 2017). In prion disease, PrP^C^ is not only necessary for the generation of PrP^Sc^ but is also involved in mediating neurotoxicity (Brandner et al., 1996). Many lines of evidence indicate that PrP^C^ is rate-limiting for the progression of prion diseases, and hemizygous *Prnp*^+/o^ mice expressing approximately 60% of wildtype PrP^C^ levels enjoy a vastly extended life expectancy after prion inoculation (Bueler et al., 1994). Therefore, it was proposed that quenching the availability of PrP^C^ may represent a feasible therapeutic strategy against prion diseases (Vallabh et al., 2020).

A number of candidate compounds binding to or lowering PrP^C^ levels have been reported (Karapetyan et al., 2013; Silber et al., 2014). However, small molecules often display pleiotropic actions and off-target effects, and compounds that lack a well-defined target do not allow drawing far-reaching conclusions about the biosynthesis of PrP^C^. Therefore, only few modulators of PrP^C^ biosynthesis, including the transcription factors sXBP1 and SP1 and the direct interactor, LRP1, were identified thus far (Bellingham et al., 2009; Dery et al., 2013; Parkyn et al., 2008; Rybner et al., 2002; Shyu et al., 2002; Vincent et al., 2009).

Here we have assessed the expression of PrP^C^ in the human CNS-derived glioblastoma cell line, U251-MG, after downregulation of each protein-coding gene by arrayed RNA interference (RNAi). The screen yielded 54 novel regulators of PrP^C^. Nine of those could be validated through an inhibitory CRISPR assay (CRISPRi), and six showed inverted-polarity PrP^C^ regulation by activating CRISPR (CRISPRa). This unbiased approach enabled the discovery of unanticipated molecular players regulating PrP^C^ expression levels. The cell-based screening and validation methodologies described here can be easily adapted to identify regulators of the expression of any other protein of interest.

## Results

### Primary screening for regulators of PrP^C^ expression

To gain insight into proteins that affect PrP^C^ expression, we performed a high-throughput whole genome RNAi screen in a human glioblastoma derived cell line, U251-MG. We chose this cell line because of its euploid karyotype and its relatively high PrP^C^ expression level. We opted to not address the M/V genotype of the cells as there is no evidence that it would influence PrP^C^ expression. U251-MG cells were successfully utilized in a high-throughput screen for miRNA regulators of PrP^C^ expression (Pease et al., 2019). We used a commercially available siRNA library consisting of 64’752 unique siRNAs targeting 21’584 annotated human genes. For the primary screening round, three distinct equimolar siRNAs targeting the same transcript were pooled into 21’584 individual wells of 62 plates (1.67 µM per siRNA) to a final concentration of 5 µM. This library was denoted as “pooled library”. Subsequently, siRNAs were reformatted into final assay destination plates to a final concentration of 5 nM using an ECHO555 acoustic dispenser.

We developed a specialized plate layout with the goal of maximizing the randomization of replicas and therefore minimizing the impact of systematic errors. Each assay was run in duplicates dispensed on two individual plates and in two distinct plate locations. We used a cell-death inducing siRNA (20 nM) to control for reduction in viability (n=22 per plate) as well as pool of non-targeting (NT) siRNAs (n=44 per plate, 5 nM)) and a pool of *PRNP*-targeting siRNAs (n=22 per plate, 5 nM). Controls were distributed in a checkerboard pattern across the plate with the aim of identifying any systematic errors deriving from gradients that may arise in the plates during protracted cell culture (Fig. 1A). Assay plates containing the siRNAs were thawed, and U251-MG cells were seeded following reverse transfection with Lipofectamine.

**Figure 1.**
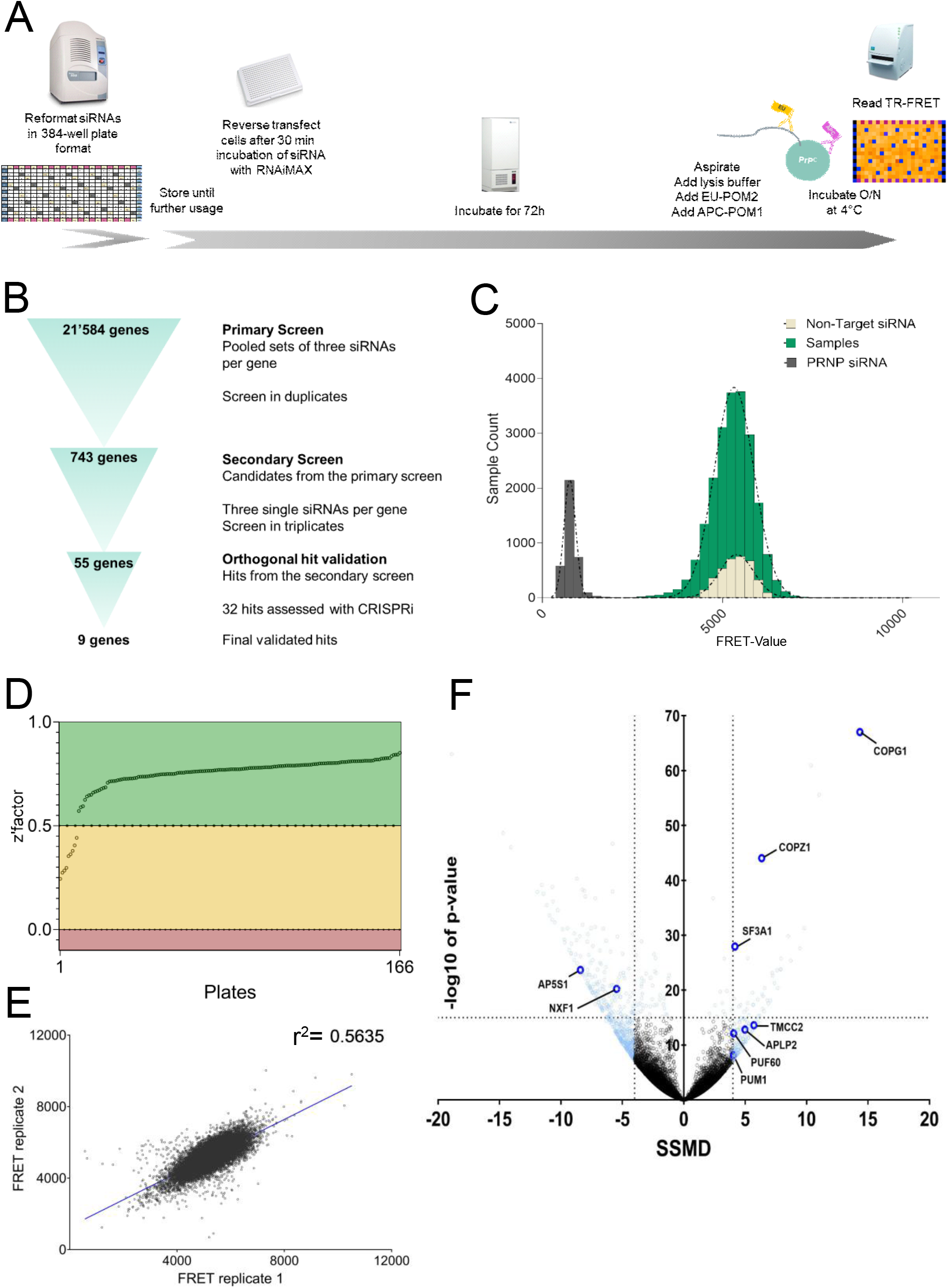
Workflow of the siRNA screen and primary screening round in U251-MG glioblastoma cell line. **(A)** Depiction of the screening workflow. **B)** Hit selection process. **C)** Distribution of populations of positive controls (*PRNP* targeting siRNAs), negative controls (non-targeting siRNAs) and samples (contents of whole genome siRNA libraries) across FRET values. The x-axis represents PrP^C^ levels measured with TR-FRET; the y-axis represents the number of assays grouped for the given FRET range. Controls showed a strong separation allowing a reliable detection of changes in PrP^C^ levels. Most of the sample population did not show a significant regulation when compared to the non-targeting controls. **D)** Z’ factor of each plate from the primary screen reporting the separability between the positive and negative controls. **E)** Duplicate correlation over all the samples from the primary screen. The Pearson correlation coefficient, r^2^-value, is depicted in the graph. **F)** Volcano plot displaying -log_10_p-value and SSMD scores across the whole genome dataset. 743 candidates identified in the primary screen are colored in light blue. Final nine hits are colored in dark blue and labelled.

After three days of culture, cells were lysed and PrP^C^ levels was measured using a solution-based immuno-assay. Two antibodies directed against non-overlapping epitopes of PrP^C^, POM2 coupled to Europium (POM2-EU) as donor and POM1 coupled to allophycocyanin (POM1-APC) as acceptor, were added to the lysates. Binding of the antibodies to PrP^C^ brings the europium and the allophycocyanin into close proximity, allowing for time-resolved fluorescence resonance energy transfer (TR-FRET) and emission of longer-wavelength light whose intensity is proportional to the concentration of PrP^C^. For quality control, we first inspected heat maps plotted for raw TR-FRET values per well after correction for spectral overlapping (Ballmer et al., 2017). We then calculated the standardized mean difference (SSMD) and Z’ factor (Zhang et al., 1999; Zhang, 2011) reporting the separation of positive (*PRNP* targeting siRNAs) and negative (NT siRNAs) controls.

We measured 166 plates of 384 wells, totaling 63’744 measurements The outermost wells did not contain any samples and were excluded from Z’ factor and SSMD calculations due their proneness for evaporation (ig 1C and D and Supplementary Fig. 1A). We considered that inhomogeneities of temperature, humidity or CO_2_ concentration during tissue culturing might create artifactual signal gradients which could impair the interpretation of the results. However, no such artifacts were observed across the whole-genome dataset. Z’ factors, which report the discriminatory power between positive and negative control, were >0.5 and 0-0.5 for 157 and 9 plates respectively. These values indicate that the screen was robust and allowed for reliable hit calling (Zhang et al., 1999). Inter-plate variability was assessed by calculating the Pearson correlation coefficient between duplicates, and yielded a value of 0.56, indicating that the screen data was sufficiently robust to select candidate genes (Fig. 1E). We used strictly standardized mean differences (SSMD) to assess the effect of each target on PrP^C^ expression (Zhang, 2011). Targets with an effect size <-4 or >4 on PrP^C^ expression were considered candidates for a secondary screen. We selected this threshold based on the weakest cumulative SSMD for controls on all plates (Supplementary Fig. 1), found to be -5, to be inclusive of all candidates of interest. As discordant duplicates occurred rarely (Fig. 1E), they were also included as candidate genes if reaching the threshold of p >10^−15^ (Student’s t-test) which is not sensitive to replica discrepancies.

In summary, 743 out of 21’584 tested genes were selected as candidates to be assessed in a secondary screen (SSMD < -4 or > 4 and/or p >10^−15^; Supplementary Table 1). When suppressed by siRNA, 563 of these reduced and 180 genes increased PrP^C^ levels (Fig. 1F). We shall henceforth refer to such genes as “stabilizers” and “limiters” of PrP^C^ expression, respectively.

### Hit validation through secondary screens

Off-target effects are a common caveat of siRNA screens, and can arise because siRNA seed sequences may display homology to illegitimate loci in the genome (Jackson et al., 2003; Jackson and Linsley, 2010). In order to address this potential problem, we performed a secondary deconvolution screen. Whilst the primary screen had been performed using mixtures of three siRNAs against each gene, in the secondary screen each candidate was subjected to the three targeting siRNAs individually in U251-MG cells. Each experiment was run in three replicates, leading to 9 assays per gene for all 743 candidate genes from the primary screen. The screen was run in two subsets of genes: a first round encompassing 583 genes (Fig. 2A, black dots) and a second round consisting of 160 genes (Fig 2A, purple dots). To add stringency and robustness to this approach, the same single-siRNA screen was performed with a second cell line, GIMEN, which was chosen for its neuroectodermal origin (Kuzyk et al., 2016) and high endogenous PrP^C^ expression (see Fig. 4B).

**Figure 2.**
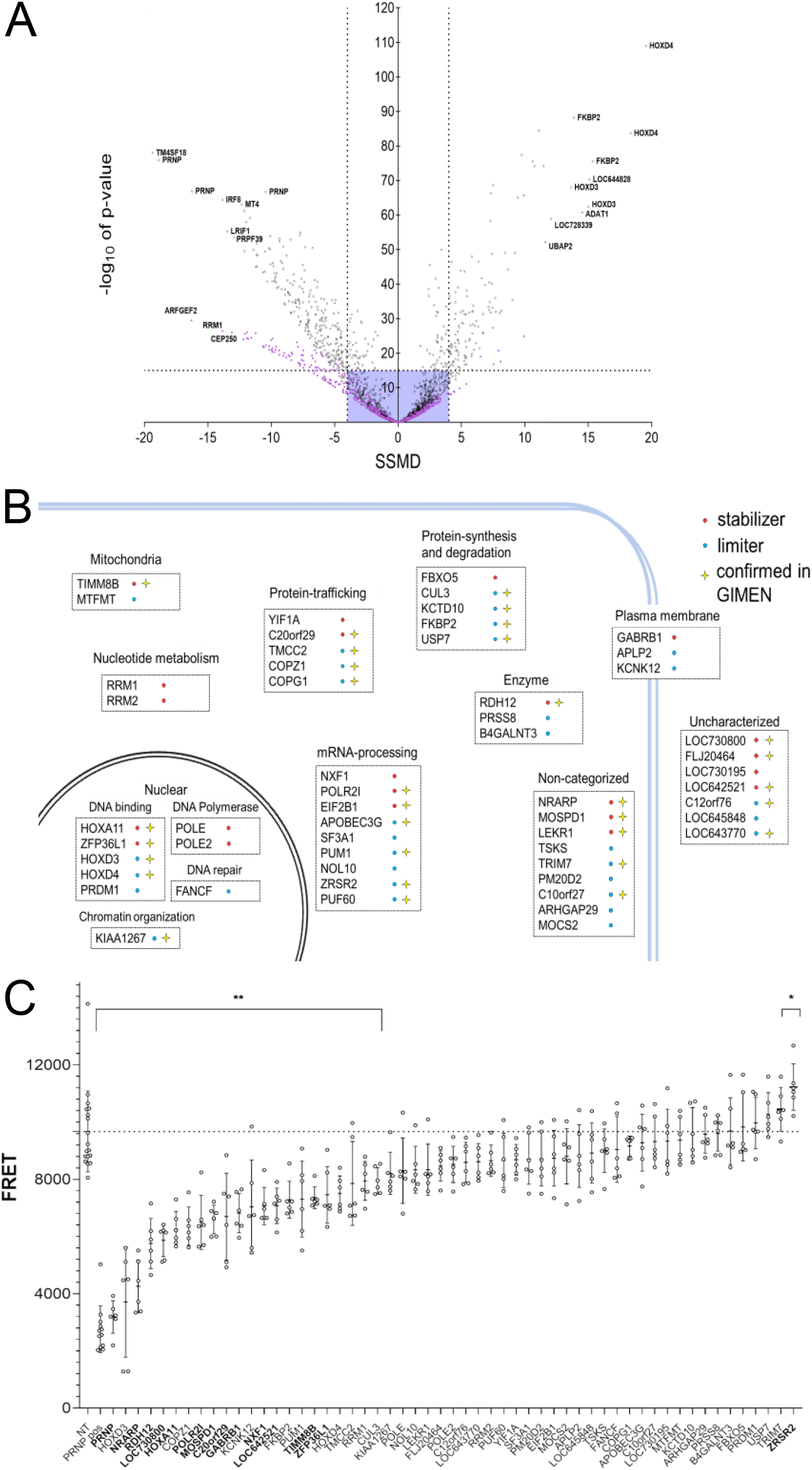
Secondary screening and validation of hits in smNPC-derived neurons. **A)** Extent (SSMD) and confidence (p-value) of PrP^C^ modulation by candidate limiters (left) and stabilizers (right). Dots represent averages of triplicates; 3 siRNAs/gene were assayed. The 10 strongest limiters and stabilizers, as well as siRNAs for *PRNP* are highlighted. Violet box: cut-off criteria. Targets with ≥2 siRNAs with |SSMD| > 4 or p < 10^−15^ were considered hits (n=54). Colors indicate screening subsets (black= first round, violet= second round). **B)** Function and topology of the top 54 hits. Red: stabilizers; limiters: blue. Star: hits overlapping with GIMEN cells. **C)** PrP^C^ levels in smNPC-derived neurons transfected with a pooled set of siRNAs (as used for the primary screen, 10 nM). Dotted line: average value for cells transfected with non-target siRNAs. Genes highlighted in bold showed the same effect observed in the siRNA screening performed in U251-MG cells. n=6 individual wells for each set of siRNAs. Values represent mean ± SD. * p = 0 0283 ** p ≥ 0 0068 (Dunnett’s multiple comparisons test)

Both single-siRNA screens were carried out similarly to the primary screen, except that each assay was run as a triplicate rather than duplicate. Hits were called if ≥2 of the 3 siRNAs had a strong effect (SSMD < -4 or SSMD > 4 or p < 10^−15^). Using these criteria, we identified 54 hits (excluding *PRNP*) in U251-MG cells (Fig. 2A, B), of which 31 were confirmed in GIMEN cells (Supplementary Table 1). Out of the 54 hits, 22 and 32 genes were classified as stabilizers and limiters, respectively.

### Validation of the hits with siRNAs in smNPC-derived neurons

PrP^C^ is highly expressed in human neurons (Bendheim et al., 1992), which are key players in prion diseases (Mallucci et al., 2007). We therefore tested the 54 hits in neurons derived *in vitro* from small molecule neural progenitor cells (smNPCs) (Dhingra et al., 2020). Firstly, we assessed the endogenous expression of PrP^C^ in smNPCs and smNPC-derived neurons at 11 days post differentiation. smNPCs and smNPC-derived neurons showed detectable levels of PrP^C^ (Supplementary Fig. 2B). We then tested the efficiency of siRNA transfection. Four-day old smNPC-derived neurons were transfected individually with three distinct *PRNP* targeting and one non-target siRNAs (10 nM). On day 11, cells were harvested and PrP^C^ levels were assessed by immunoblotting. The three siRNAs showed variable effects on PrP^C^ suppression (Supplementary Fig. 2C) similarly to the secondary screening, where *PRNP* siRNA 3 also showed the weakest effect.

We then investigated whether the 54 hits identified in the secondary screen produced similar effect in smNPC-derived neurons. We opted to test the effect of each gene by pooling three siRNAs, analogous to the primary whole-genome screen. siRNAs targeting the 54 hits, *PRNP-targeting* siRNAs and non-targeting siRNA controls were reformatted in a 96-well plate. smNPC-derived neurons were seeded in a separate 96-well plate and cells were transfected with siRNAs after 4 days of differentiation.

Seven days later, cells were lysed, and lysates were applied to TR-FRET readout (Fig. 2C). We found that 14 targets showed a conserved effect in smNPC-derived neurons. Of these, 13 were stabilizers and one was a limiter.

### CRISPRi validation of hits in dCas9-KRAB U251-MG cells

In order to increase confidence in the hits identified by the secondary screening and to eliminate false-positives due to off-target effects of the siRNAs, we performed CRISPRi experiments in U251-MG cells. We chose CRISPRi as an additional method, as it acts through a repressor domain (KRAB) fused to the Cas9 protein, therefore achieving gene regulation through inhibition of transcription (Qi et al., 2013). Since single clones of dCas9-KRAB-expressing cells can show differential basal transcriptomic signatures (Stojic et al., 2018), we generated a polyclonal bulk of U251-MG^dcas9-KRAB^ cells through lentiviral transduction and kept it under blasticidin selection (10 μg/mL). Next, we constructed plasmids containing four non-overlapping guide RNAs (gRNAs) per gene of interest to achieve maximum repression. Each individual gRNA was controlled by a different housekeeping promoter (hU6, mU6, hH1 and hS7K). For control, we used two sets of each 4 gRNAs bearing non-targeting sequences with no known homology to any mammalian genes. Positive control contained a construct with 4 gRNAs against *PRNP* (Supplementary Fig. 2A). A total of 32 genes were further assessed with CRISPRi. We opted to exclude 22 candidates from further analysis based on low abundance of transcripts in the U251-MG line or known global regulators of protein production (expression data summary can be found in Supplementary Table 1). We seeded 2×10^5^ U251-MG^dCas9-KRAB^ cells in 6-well plates and transduced them on the next day with lentiviruses containing gRNAs for each of the candidate genes. At 3 days post transduction, we selected the transduced cells with puromycin (1 μg/mL) for 5 further days (Supplementary Fig. 2B). Lysates were then subjected to either TR-FRET to assess PrP^C^ levels or quantitative real-time PCR (qRT-PCR) to determine the efficiency of the CRISPRi treatment (list of all primers used in the study can be found in Supplementary Table 1). CRISPRi yielded efficient downregulation of 27 candidates after 5 days of selection (Fig. 3A and Fig. 3B, also see Supplementary Fig. 2C, for two genes ran independently). *NXF1*, a nuclear export factor involved in mRNA export (Chen et al., 2019) did not yield an apparent downregulation potentially due to its interference with the house keeping gene, however, PrP^C^ levels were altered. Ten of the 32 tested hits were found to regulate PrP^C^ in CRISPRi experiments (Fig. 3A and Supplementary. Fig. 2C).

**Figure 3.**
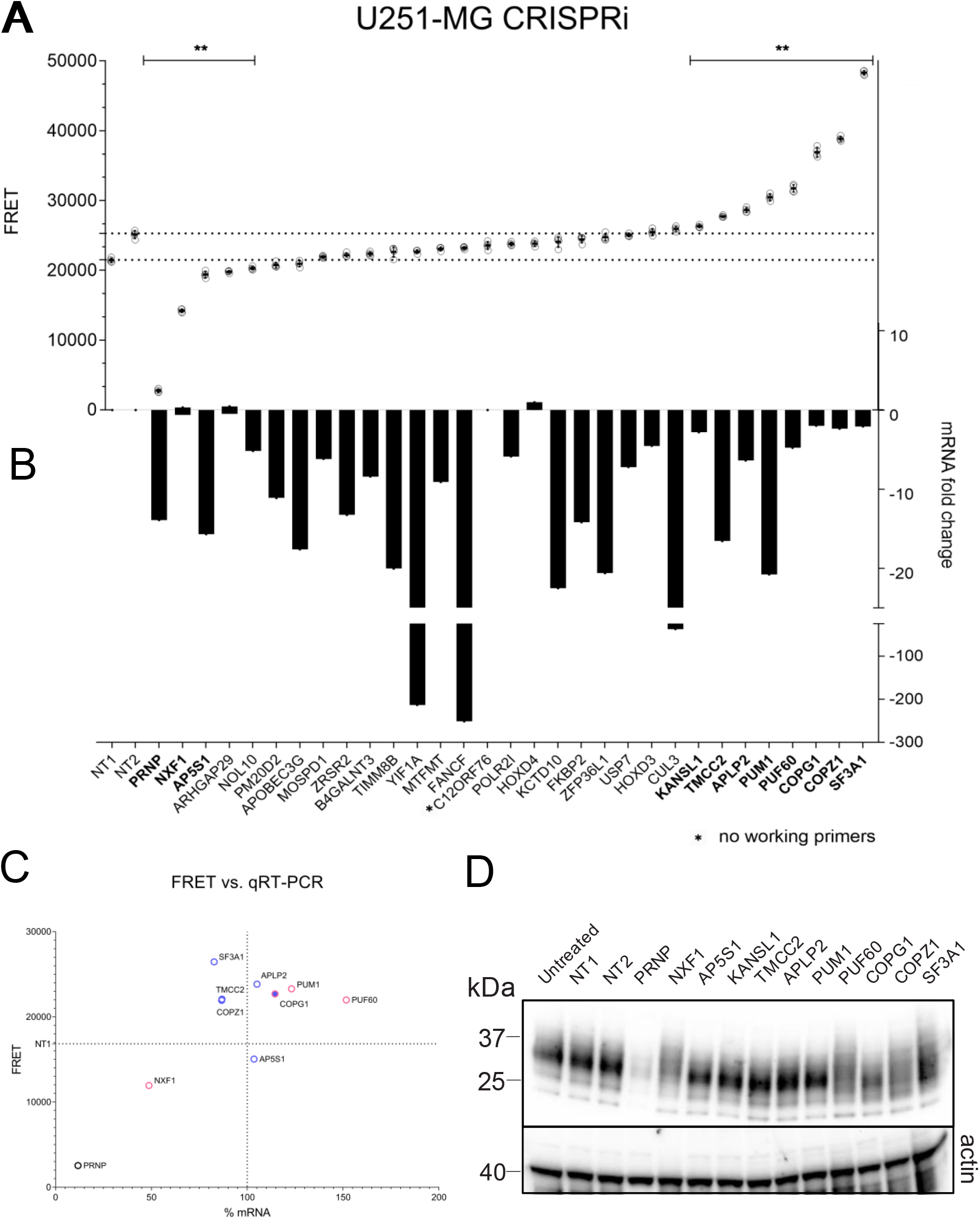
Hit validation in U251-MG cells by CRISPR interference. **A)** PrP^C^ protein levels of dCas9KRAB-U251-MG cells transduced with gRNA CRISPRi lentiviruses against the targets depicted in B. Mean values ± SD (n=4 technical replicates) are shown ** p ≥ 0 0023 (Dunnett’s multiple comparisons test) **B)** CRISPRi activity measured by the mRNA level of the target gene in comparison to a negative control sample (NT1) after normalization to the housekeeping gene *ACTB. C12orf76* could not be tested due to non-functional primers. Three targets showed an increase in mRNA levels, and 27 genes showed the expected decrease in mRNA levels. Two constructs bearing non-targeting gRNAs were used for control. Only genes which showed a statistically significant difference to both NTs were considered to be true hits. Hits with same directionality and effect obtained in the siRNA screens are highlighted in bold. Bars represent averages of three replicates. **C)** Modulation of *PRNP* mRNA and PrP^C^ protein levels after CRISPRi mediated suppression of 2 stabilizers and 7 limiters. qRT-PCR and FRET analysis of *PRNP* mRNA of the nine hits with a significant effect on PrP^C^ levels, in an independent repetition experiment. Normalization to *ACTB*. Three genes (pink) regulated *PRNP* mRNA levels, whereas five (blue) modulated PrP^C^ levels post-transcriptionally and one gene (pink circle with blue filling) acted both transcriptionally and post-transcriptionally. **D)** Western blot analysis of the same 10 samples as in B, probed with antibody POM2 against PrP^C^. The FRET and immunoblot results were congruent. In addition, NXF1, PUF60, COPZ1, COPG1 and SF3A1 induced a shift in the glycosylation pattern of PrP^C^.

**Figure 4.**
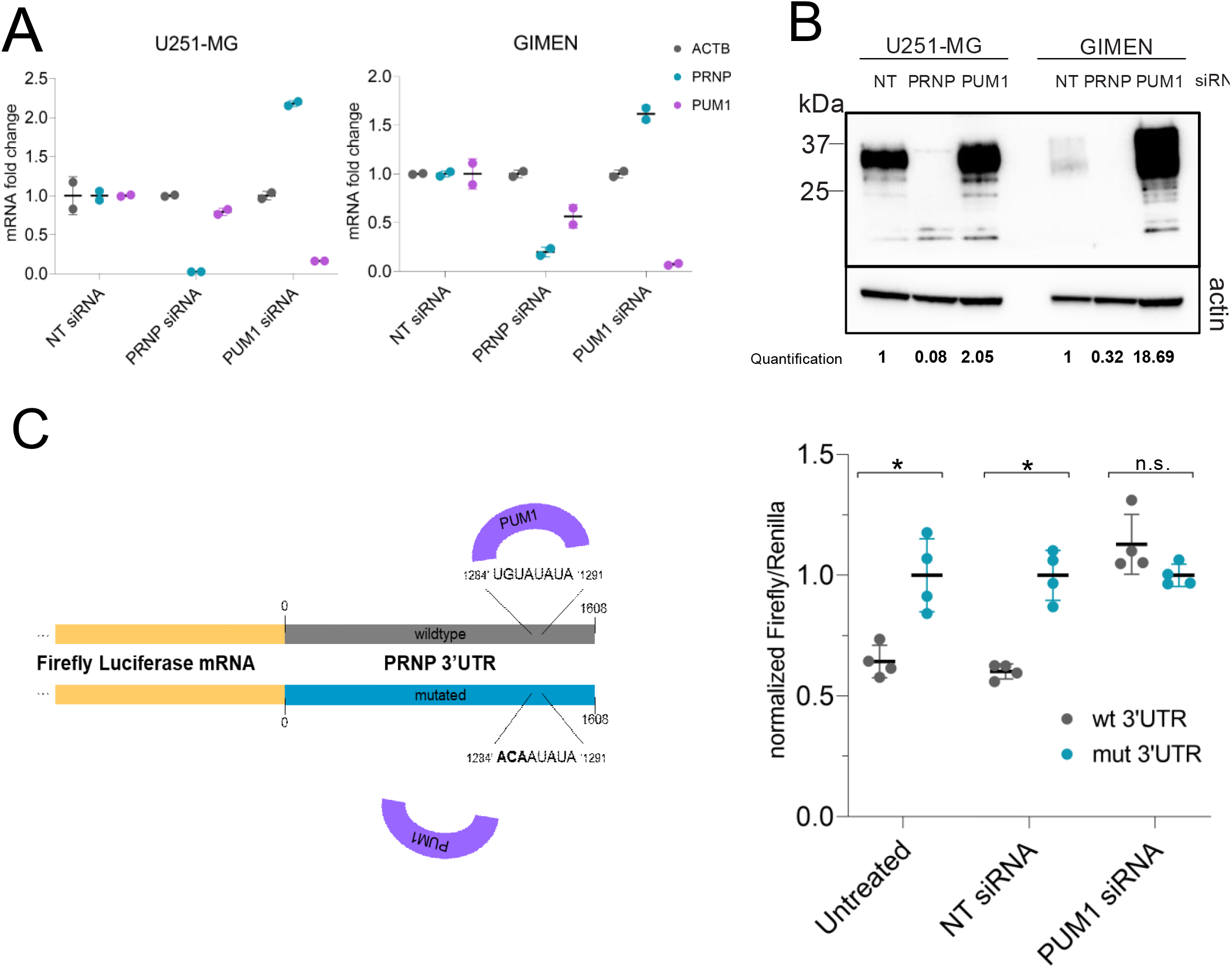
PUM1 regulates PRNP mRNA via its 3’UTR. **A)** qRT-PCR analysis of U251-MG and GIMEN cell lines upon transfection with *PUM1* siRNA in 6-well format. *PUM1* mRNA was efficiently downregulated Δ t values were normalized to those of β-actin (*ACTB*). Downregulation of *PUM1* resulted in increased *PRNP* mRNA in both cell lines. *PRNP* siRNA was used for control. **B)** Western blot analysis of U251-MG and GIMEN cell lines upon transfection with *PUM1* siRNA. *PUM1* downregulation lead to an upregulation of PrP^C^ levels (as seen in Fig. 3B through CRISPRi). Antibody POM2 was used for detection of PrP^C^. *PRNP* siRNA was used for control. For quantification, signal intensity of PrP^C^ was normali ed to the signal intensity of β-actin **C)** Dual-Glo Luciferase assay to assess regulation of *PRNP* mRNA by *PUM1* via its 3’-UTR. Left panel shows a schematic of the assay. The numbers indicate the position in the 3’-UTR of *PRNP*. The *PRNP* 3’-UTR sequence, predicted to bear the consensus sequence for *PUM1* binding, was placed after the gene coding for the firefly luciferase. Through a mutation in the 3’-UTR binding site of *PUM1* (wt 3’-UTR => mut 3’-UTR), the binding is prevented leading to increased expression of the firefly luciferase. Two plasmids (wt 3’-UTR and mut 3’-UTR) were co-transfected into HEK293-T cells either with none, NT, or *PUM1* siRNAs. Cells transfected with the mut 3’-UTR plasmid showed a higher signal in the assay in comparison to transfection with the wt 3’U R plasmid Similar results were obtained for co-transfection of mut 3’-UTR plasmid and a non-targeting siRNA. The co-transfection with *PUM1* siRNA abrogated the signal difference. Means ± SD (n=4). * p ≤ 0.016, n.s. = non-significant (multiple t-test).

### Mechanism of action of PrP^C^ regulators

The hits described here may regulate PrP^C^ by affecting its transcription, its translation, or its turnover. In order to distinguish between these scenarios, we challenged the ten hits with CRISPRi, to additionally assess mRNA levels of *PRNP* using qRT-PCR, in combination with PrP^C^ levels by western blotting and TR-FRET. KANSL1 did not reach statistical significance and was excluded from further analyses. This final selection round yielded APLP2, TMCC2, C20orf29/AP5S1, SF3A1, COPZ1 as post-translational regulators of PrP^C^. COPG1 was found to change PrP^C^ levels transcriptionally and post-translationally. Instead, PUM1, PUF60 and NXF1 were found to regulate PrP^C^ by altering *PRNP* mRNA levels (Fig. 3C, Supplementary Fig. 2D). In addition, the band patterns on Western blots indicate that CRISPR interference with NXF1, PUF60, COPZ1, COPG1 and SF3A1 affected the glycosylation of PrP^C^ (Fig. 3D). We conclude that CRISPRi induced highly efficient and selective repression of select genes, thereby representing a valid tool for the independent confirmation of the results obtained from RNAi experiments.

### Role for genes that regulate PrP^C^ expression in sporadic Creutzfeldt-Jakob disease (sCJD) susceptibility

We tested for a role of genes identified in the primary screen (n=743) or after hit validation (n=9) in genetic susceptibility to sCJD using data from a recent collaborative genome-wide association study (Jones et al., 2020). Gene based tests, such as those implemented with MAGMA and VEGAS packages (de Leeuw et al., 2015; Mishra and Macgregor, 2015), aggregate single nucleotide variants within entire genes into a single statistical test, whilst accounting for linkage disequilibrium and other confounders like gene size. No genes surpassed thresholds that consider multiple testing (Supplementary Table 1). The top ranked association from the primary screen was *MPPED1* (MAGMA unadjusted p=0.0017), and from the validated hits, *SF3A1* (MAGMA unadjusted p=0.024).

### CRISPRa validation of hits in dCas9-VPR U251-MG cells

To assess whether an opposing regulation could be achieved through the activation of the final 9 hits, we performed CRISPRa. In contrast to a repressor domain used in CRISPRi experiments, the endonuclease deficient Cas9 (dCas9) in this instance is coupled to a transcriptional activator domain, VP64-p65-RTA (VPR), leading to efficient activation of target genes (Chavez et al., 2015). In a similar setup to the CRISPRi experiments, we generated a polyclonal bulk of U251-MG^dCas9-VPR^ cells and later transduced these cells using four gRNAs per gene targeting the nine hits. We included *PRNP* targeting control gRNAs as well as scrambled non-targeting gRNA sequences.

Differentially to the CRISPRi experiments, CRISPRa led to an efficient upregulation of *PRNP* on protein and mRNA levels already at three days post-selection (Supplementary Fig. 3A, 3B and 3C). All 9 hits were efficiently upregulated in mRNA levels as measured by qRT-PCR (Supplementary Fig. 3D) with at least one construct targeting different TSS. TMCC2, APLP2, PUM1, COPG1, COPZ1 and SF3A1 led to regulation of PrP^C^ measured with TR-FRET and visualized with immunoblotting in the opposing direction to the CRISPRi results (Supplementary Fig. 3E, 3F), suggesting these 6 hits have a bidirectional effect on PrP^C^ levels.

### PUM1 mediates decay of PRNP through binding its 3`UTR

One of the 9 hits was Pumilio-1 (PUM1), an RNA-binding protein that binds its targets through their 3’-untranslated Region (3’-UTR) and mediates their decay (Van Etten et al., 2012). PUM1 binds its targets through a well-established consensus sequence, the Pumilio responsive element (PRE, 5’-UGUANAUA-3’) (Bohn et al., 2018). We first tested whether the PRE was present on the 3`UTR of *PRNP*. Using the RPI-Seq tool (Muppirala et al., 2011), we identified a PRE within the 3`UTR of *PRNP*. To confirm that PUM1 is a regulator of *PRNP* mRNA, we performed independent siRNA transfections in 6-well plates in both cell lines, U251-MG and GIMEN. We then subjected the extracted RNA to real-time quantitative PCR (qRT-PCR) and cell lysates to immunoblotting. In both cell lines tested, we confirmed that PUM1 silencing had an upregulating effect on *PRNP* mRNA, which also resulted in higher protein levels (Fig. 4A-B).

We then used a dual-luciferase reporter assay to test whether the effect observed was due to the predicted interaction of PUM1 with the 3’-UTR of *PRNP* mRNA. The wild-type (wt) and a mutated (mut) version of the *PRNP* 3’-UTR were cloned into pmirGLO vectors encoding Renilla and Firefly luciferase. These two enzymes report transfection efficiency and activity of the inserted construct, respectively. The mutation induced spanned three base pairs (bp) of the PRE on *PRNP* 3’-UTR to impair the binding of PUM1 as previously reported (Kedde et al., 2010). Plasmids were then transfected into HEK293T cells in presence or absence of either NT or PUM1 targeting siRNAs. Luminescence arising from the Firefly luciferase was measured after 48 hours and was followed by the measurement of Renilla luciferase for normalization. In the absence of siRNAs as well as in presence of NT siRNA treatment the mutated 3’-UTR containing construct yielded a higher signal in comparison to the wt 3’-UTR containing construct. When cells were treated with PUM1 targeting siRNAs the difference seen in the decrease in signal for the wild-type construct was abolished, indicating that PUM1 is indeed acting on the 3’-UTR of *PRNP* mRNA (Fig. 4C).

## Discussion

The concentration of PrP^C^ in any given cell type depends on an equilibrium between its biosynthesis and degradation rates, which in turn are controlled by a multitude of processes – from mRNA transcription to cotranslational secretion into the endoplasmic reticulum, quality control of folding, glycolipid bonding, vesicular transport, and eventually proteolysis as well as extracellular shedding. Most factors controlling these steps are unknown, and it is difficult to imagine that they could be discovered through educated guesses. Conversely, an unbiased interrogation of the entire human genome would yield PrP^C^ regulators that may not have been predicted by existing knowledge.

As *PRNP* is highly expressed in the CNS (Bendheim et al., 1992), we chose a glioblastoma cell line U251-MG for the initial round of screening, additionally due to the suitability of this cell line in high-throughput screenings (Pease et al., 2019). Our strategy relied on recursive screening rounds leading to progressive attrition of candidates. The primary screening campaign, in which each gene was targeted with a mixture of 3 siRNAs, identified 743 presumptive candidate genes strongly influencing PrP^C^ levels. However, a secondary screen with individual siRNAs showed that many of these hits could only be confirmed with one or two of the three siRNAs present in the original mixtures. These discrepancies may arise because the efficacy of suppression by the individual siRNAs was variable, or because some siRNAs exerted spurious off-target effects. We therefore decided to perform tertiary screens on those candidates that showed an effect with at least 2 siRNAs in two different cell lines. This filtering strategy represents a trade-off between the elimination of false-positive signals and the retention of assay sensitivity.

The resulting list of 54 genes was then interrogated further to assess their effect on human neurons differentiated in vitro. These investigations confirmed most of the hits identified as PrP^C^ stabilizers, but only one limiter gene. The bias towards stabilizers may be a consequence of the long half-life of proteins in post-mitotic neurons (Dörrbaum et al., 2018) which may conceal the effects of short RNAi regimens, and suggests that most stabilizers may directly affect *PRNP* biosynthesis. We addressed these questions by suppressing the genes of interest with an independent methodology. CRISPRi has recently emerged as a potent tool to reliably control gene expression (Gilbert et al., 2013; Qi et al., 2013). We used multiple gRNA sequences per target, with the intent to increase the efficiency of modulation (Kurata et al., 2018). Through CRISPRi we identified nine hits significantly regulating PrP^C^, six of which controlled PrP^C^ levels post-translationally (Supplementary Table 1). The partial discordance between the results obtained through siRNAs and CRISPRi is not unexpected. Firstly, the mode-of action of the methods chosen are distinct from each other. In our setup, we applied siRNAs for a duration of three days, which leads to an acute depletion of the target mRNAs, however, to achieve efficient CRISPRi, the cells were cultured for eight days, during which compensatory mechanisms controlling gene expression may take place. Secondly, the use of two non-targeting controls for CRISPRi as opposed to one non-targeting control for siRNAs increases the stringency for calling hits. Thirdly, potential off-target effects seen through one of the methods can be distinct from the other.

Because of the reasons expounded above, the nine genes identified in both ap proaches represent, with a high degree of confidence, true regulators of PrP^C^. To explore if the regulation of PrP^C^ by the final hits occurs bidirectionally, we made use of a second system, CRISPRa, relying on an endonuclease deficient Cas9 coupled to a transcriptional activator domain consisting of VP64-p65-RTA (VPR) (Chavez et al., 2015). We found that 6 of the 9 hits acted in an opposing direction on PrP^C^ levels, upon their activation with CRISPRa. One of the latter hits, Pumilio 1 (PUM1), is a known mediator of degradation of transcripts through binding to the 3’U R on their mRNA (Van Etten et al., 2012). *PRNP* was not identified in a previous study of PUM1-regulated genes (Bohn et al., 2018), perhaps because of insufficient levels of PrP^C^ expression in HEK cells. Sequence inspections then revealed the PUM1-binding consensus sequence in the human, but not mouse, *Prnp* 3’U R, and mutational analyses of the consensus sequence confirmed the mechanism of this regulation. Besides clarifying the molecular mechanism by which PUM1 modulates PrP^C^, these data provide direct evidence for the validity of the regulators identified in our screen.

We failed to identify any transcription factors specifically controlling PrP^C^ expression in the CNS, perhaps because transcriptional gene regulation relies on redundant factors. Also, the cancer cell line used for the primary screen, while derived from a CNS tumor, may not be representative of transcriptional regulation within the CNS.

Some of the regulators identified in this screen were not entirely unexpected. The members of the COPI complex, COPG1 and COPZ1, are involved in the retrograde transport of vesicles from the Golgi apparatus to the ER. Interference with the COPI complex is known to affect the expression of membrane proteins such as APP (Bettayeb et al., 2016). Furthermore, AP5S1 is part of the newly described Fifth Adaptor Protein Complex (AP-5) which recycles proteins out of late endosomes (Hirst et al., 2011). The decrease of PrP^C^ after AP5S1 knock-down may be due to decreased retrieval from late endosomes.

Other regulators were entirely unexpected, and their mechanism of action is not obvious. The ablation of APLP2 had no significant effect on incubation times of scrapie in mice (Tamgüney et al., 2008), suggesting that the upregulation of PrP^C^ upon APLP2 knockdown may be specific to humans. Mechanistically, APLP2 might serve as a co-receptor for the endocytosis of PrP^C^, akin to what has been described for MHC class I molecules (Peters et al., 2013) Alternatively, the regulation of PrP^C^ may be related to the role of APLP2 in metal homeostasis (Millhauser, 2004; Roisman et al., 2019). TMCC2 has been previously described as an ER-resident molecule that interacts with APP and has an effect on metabolism of amyloid-β (Hopkins et al., 2011).

In view of the analogies between prion diseases and Al heimer’s disease (Aguzzi and Haass, 2003), the regulation of PrP^C^ via TMCC2 may occur through a common mechanism. NXF1 and PUF60, well known RNA-regulating proteins, may directly influence the availability of *PRNP* mRNA. Finally, in the case of SF3A1, a splicing factor, the regulation of PrP^C^ may be indirect as there are no known mechanisms by which SF3A1 could interfere with the formation of PrP^C^ at the protein level. To extend the understanding of the regulatory network of PrP^C^ expression, we also cross-referenced the nine hits to miRNAs hits previously identified by our group (Pease et al., 2019). The study identified four miRNA targets for which no direct interaction with *PRNP* mRNA was established. However, based on the target prediction data, miRDB, (Chen and Wang, 2020) we did not find any plausible links between the identified miRNA hits and the nine hits reported here.

In addition to the mechanisms discussed above, the biogenesis of PrP^C^ involves co-translational secretion of the nascent polypeptide chain into the lumen of the ER and addition of a glycophosphoinositol (GPI) anchor. However, not all of the players known to control these processes were identified as hits. For one thing, some of these genes encode essential proteins whose knockdown may drastically decrease cell viability. Furthermore, because the pathways involved in proteostasis are multiple and partially redundant (Andréasson et al., 2019), suppressing a single gene may not result in a measurable effect on PrP^C^ levels.

In many neurodegenerative diseases including Al heimer’s and ar inson’s disease fundamental clues to the pathogenesis were provided by the study of families with Mendelian disease transmission. However, in the case of prion diseases these strategies have been largely unsuccessful. Although genome-wide association studies (GWAS) have identified two potential risk loci (Jones et al., 2020), the only strong genetic risk factor for Creutzfeldt-Jakob disease has remained *PRNP* itself (Hsiao et al., 1989). This sobering situation was a primary driver of the present study, as the assessment of every individual protein-coding genes may plausibly highlight factors undetectable by human genetics. In addition, we did not find evidence of a genetic association between screen hits and sCJD susceptibility using gene-based analysis from a recent GWAS (Jones et al., 2020). Similarly, in the GWAS, PrP brain expression quantitative trait loci near to *PRNP* showed no evidence of association with sCJD. Expression level of PrP is a powerful determinant of incubation time in rodent models of prion disease (Büeler et al., 1993), but overexpression does not appear to effect susceptibility to infection (Douet et al., 2014). Whether the genetic regulators of PrP expression in cells determine aspects of the phenotype of the human disease such as age at onset or clinical duration remains to be determined. The ranked genetic associations of the screen hits made available here may be useful in prioritization of genes for future studies. The modifiers enumerated here provide unexpected insights into a vast and highly diverse regulatory network of PrP^C^, and may eventually provide druggable targets for development of therapeutics in addition to the direct suppression of PrP^C^ (Minikel et al., 2020; Raymond et al., 2019), for instance through mRNA-based therapeutics to modulate PrP^C^-limiters (Martini and Guey, 2019).

## Material and Methods

### Cell culturing

U-251MG (Kerafast, Inc., Boston, MA, USA, AccessionID: CVCL_0021) and GIMEN (CLS Gmbh, Eppelheim, Germany AccessionID: CVCL_1232) cells were cultured in T150 tissue culture flasks (TPP, Trasadingen, Switzerland) in OptiMEM without Phenol (Gibco, Thermo Fisher Scientific, Waltham, MA, USA) supplemented with 10%FBS (Takara, Göteborg, Sweden), 1% NEAA (Gibco), 1% GlutaMax (Gibco), and 1% Penicilin/Streptomycin (P/S) (Gibco). HEK293-T cells (AccessionID: CRL_3216) were cultured DMEM without phenol (Gibco) with the same conditions for aforementioned supplements. In preparation for the screening, cells were expanded, either harvested using Trypsin-EDTA 0.025% (Gibco) or Accutase (Gibco), washed with PBS (Kantonsapotheke, Zurich, Switzerland) and resuspended in Penicilin/Streptomycin free medium, pooled, and counted using TC20 (BioRad) Cell Counter with trypan blue (Gibco). Culturing of the cells for the smNPC-derived neurons was done using a published protocol (Dhingra et al., 2020).

### siRNA library preparation and reformatting

Whole genome Silencer Select Human Genome siRNA Library V4 (Thermo Fisher Scientific), which includes three individual siRNAs targeting each gene (64752 total siRNAs targeting 21.584 human genes) was purchased at the quantity of 0.25 nmol. Upon delivery, the lyophilized library was resuspended in RNAse free water (Thermo Fisher Scientific) to a final concentration of 5 µM. siRNAs targeting the same gene were aliquoted in a single transfer step using ViaFlo equipment (Integra, Konstanz, Germany) either as pooled siRNAs (referred to as the pooled library) targeting the same gene or as single siRNAs (referred to as the single library) into ECHO acoustic dispenser (Labcyte, San Jose, CA, USA) compatible 384-well LDV plates (Labcyte). For the primary screen, pooled siRNAs and control siRNAs were dispensed in duplicates at a final concentration of 5nM (20 nM for Cell Death control, Qiagen, Hilden, Germany) into white 384-well CulturPlates (Perkin Elmer, Beaconsfield, UK) using an ECHO 555 acoustic dispenser according to an optimized plate layout (Pease et al., 2019) and stored at -40°C until further use. For the secondary screen, individual siRNAs were dispensed in triplicates. As controls, a scrambled non-targeting siRNA (NT) (Ambion, Thermo Fisher Scientific) as negative and a PRNP targeting siRNA (Ambion, Thermo Fisher Scientific) as positive control, were reformatted to destination plates either as three siRNAs in pooled format or as single siRNAs. All siRNA sequences are accessible under the PubChem repository, NCBI (PubChem, 2013).

### Screening workflow

Cell culture plates containing reformatted siRNAs were thawed and 5 µL RNAiMAX (Invitrogen, Carlsbad, CA, USA) (1.8% v/v, final conc. 0.3%) diluted in P/S free medium was dispensed using a multi-drop dispenser (MultiFlo FX, Biotek, Vinooski, VT, USA). Plates were centrifuged at 1000xG for 1 minute (Eppendorf 5804R, Hamburg, Germany) and incubated for 30 min at room temperature (RT) Afterwards 6’000 cells (U251-MG) or 8’000 cells (GI-ME-N) per well were seeded in 25 µL P/S free medium and plates were incubated in a rotating tower incubator (LiCONiC StoreX STX, Schaanwald, Liechtenstein). For the primary screen, plates were removed from the incubator after 70 hours and 10 µL of 4x RT-Glo substrate and enzyme per well (Promega, Madison, WI, USA) diluted in P/S free medium, was added. Plates were incubated for another 2 hours in the incubator. Prior to measuring luminescence temperature of EnVision plate reader (Perkin Elmer) was set to 37°C and luminescence was measured without disturbances in temperature. Subsequently, medium was removed by inverting the plates, and cells were lysed in 10 µL lysis buffer (0.5% Na-Deoxycholate (Sigma Aldrich St. Louis, MO, USA) 0.5% Triton X (Sigma Aldrich), supplemented with EDTA-free cOmplete Mini Protease Inhibitors (Roche, Basel, Switzerland) and 0.5% BSA (Merck, Darmstadt, Germany). Following lysis, assay plates were incubated on a plate shaker (Eppendorf ThermoMixer Comfort) for 10 min (4°C, 700 rpm shaking conditions) prior to centrifugation at 1000xG for 1 min and incubated at 4°C for two additional hours. Following incubation, plates were centrifuged once more under same conditions mentioned above and 5 µL of each FRET antibody pair was added (2.5 nM final concentration for donor and 5 nM for acceptor, diluted in 1x Lance buffer, (Perkin Elmer)). For FRET, two distinct anti-PrP antibodies, POM1 and POM2 (Polymenidou et al., 2008), targeting different epitopes of PrP^C^ were coupled to a FRET donor, Europium (EU) and a FRET acceptor, Allophycocyanin (APC), respectively, following previously reported protocols (Ballmer et al., 2017). For APC – POM1 coupling, Lightning-Link APC Labeling Kit (Lucerna Chem, Lucerne, Switzerland) was used following manufacturer`s instructions. Plates were centrifuged once more and incubated overnight at 4°C. TR-FRET measurements were read out using previously reported parameters (Ballmer et al., 2017) on an EnVision multimode plate reader (Perkin Elmer).

### Screening data analysis

Screening data was analyzed using an in-house developed, open-source, Phyton-based high throughput screen (HTS) analysis pipeline (all documentation and code available under: https://github.com/elkeschaper/hts). Net-FRET data was calculated (Ballmer et al., 2017) and subjected to various quality control checkpoints. Initially, a heat map of individual plates was plotted to examine temperature-induced gradients or dispensing errors Subsequently ‘-Factor and strictly standardized mean difference (SSMD) scores (Zhang et al., 1999; Zhang, 2011) were calculated to report the robustness of the screens. Additionally, Net-FRET values were plotted to check for row or column effects as well as assessing correlation of duplicates or triplicates. After assessing quality of each individual plate, candidate genes were selected with the following cut-off criteria: SSMD of < -4 and > 4 *or* a p-value (t-test) of below 10^−15^. After the secondary screen, same cut-off criteria were used with the additional requirement at least 2 out of 3 individual siRNAs targeting the same transcript passing the threshold. Graphs were generated with GraphPad Prism.

### PUM1 validation and reporter assay

U251-MG or GI-ME-N cells were seeded into 6-well cell culture plates in a total culture volume of 1.5 mL. Next days, cells were transfected with NT or PUM1 siRNAs (Thermo Fisher Scientific) to a final concentration of 5 nM in a total culture volume of 2 mL. 72 hours post-transfections cells were washed once in PBS to remove cell debris and collected for immunoblotting and quantitative real-time polymerase chain reaction (qRT-PCR). For immunoblottings, cells were scraped with 100 µL lysis buffer (50 mM Tris-HCl pH 8, 150 mM NaCl, 0.5% sodium deoxycholate, 0.5% Triton-X 100, Sigma) supplemented with EDTA-FREE cOmplete Mini protease inhibitor cocktail (Sigma), incubated on ice for 20 minutes and centrifuged at 10.000xG for 10 minutes for isolation of proteins. Supernatants were then subjected to bicinchoninic acid assay (BCA) (Pierce, Waltham, MA USA) to measure total protein concentrations according to manufacturer’s instructions 25 µg of total proteins of all samples were loaded onto a 4-12% gradient gel (Invitrogen) and blotted onto a PVDF membrane (Invitrogen). Following blocking in 5% SureBlock reagent (LubioScience, Zurich, Switzerland) diluted in PBS-Tween20 (PBST, Sigma) for 30 minutes, anti-PrP antibody POM2 (Polymenidou et al., 2008) was applied to the membrane at a final concentration of 300 ng/mL in 1% SureBlock in PBST and incubated overnight at 4°C. As a detection antibody anti-Mouse HRP (BioRad) was used diluted 1:10.000 in 1% SureBlock containing PBST. Immunoblots were developed with Crescendo HRP substrate (Millipore, Dachstein, France). As loading control, an anti-Actin antibody m25 (Merck) was used at a dilution of 1:10.000 in 1%Sureblock containing PBST. Imaging was performed on Vilber (Eberhardzell, Germany) systems. Quantification was done with ImageLab (BioRad) For qRT-PCR, cells were lysed in 350 µL RLT lysis buffer (Qiagen) and RNA was extracted using RNeasy Mini kit (Qiagen) manufacturer’s instructions Concentration and quality of RNA was assessed using a NanoDrop spectrophotometer (Thermo Fisher Scientific). cDNA synthesis was done following the manufacturer’s instructions using the Quantitect Reverse Transcription kit (Qiagen). A total of 25 ng of cDNA per sample was manually transferred into 384-well PCR plates (Thermo Fisher Scientific) and SYBR green (Roche) mastermix was used for detection. Readout was performed with ViiA7 Real Time PCR systems (Thermo Fisher Scientific). As internal controls, three housekeeping genes (ACTB, TBP, GUSB) were measured for each sample. Data was analyzed and visualized using GraphPad Prism.

Wild-type 3’U R of PRNP (1608 base pairs (bp)) and a mutant version thereof, were ordered as gene blocks (Integrated DNA Technologies (IDT), Newark, NJ, USA). The mutation was positioned at the PUM1 binding site 5’-TGTATATA-3’ where was exchanged to A A in line with previous reports (Kedde et al., 2010). Additionally, the sequence was modified to contain an overlap of 25 bp at the 5’ end and 21 bp at the 3’ end with the pmirGLO Dual-Luciferase miRNA Target Expression Vector for molecular cloning into pmirGLO Dual-Luciferase miRNA Target Expression Vector (Promega). The pmirGLO vector was initially digested with PmeI (New England Biolabs (NEB), Ipswitch, MA, USA) and SalI (NEB). Either the wild-type or the mutant version of the *PRNP* 3’U R were inserted into the digested pmirGLO vector by performing a Gibson Assembly following manufacturer`s instructions (NEB). After transformation of the cloned vectors and purification of the DNA using EndoFree Plasmid Maxi Kit (Qiagen), the purified products were sent to Microsynth AG (Balgach, Switzerland) for Sanger sequencing and aligned to reference sequence of 3`UTR of PRNP using Nucleotide-Blast (NCBI). Final constructs were designated as wild-type (wt)-3`UTR-pmirGLO or mutant (mut)-3`UTR-pmirGLO. 1×10^4^ HEK-293T cells were seeded in culture medium without antibiotics in white 96-well culture plates (Perkin Elmer) prior to transfection. 100 ng of the wt/mut-3’U R-pmirGLO plasmid and 1 pmol of PUM1 or NT siRNAs (Thermo Fisher Scientific) were co-transfected in quadruplicates for each condition using 1% Lipofectamine 2000 (Thermo Fisher Scientific). Four additional wells contained no siRNAs. 48 hours post-transfection, the Dual-Glo Luciferase Reagent (Promega) was added to each well and the luminescence of the firefly luciferase was measured using EnVision plate reader (Perkin Elmer) after 20 min incubation at RT. Subsequently, Dual-Glo Stop & Glo Reagent diluted (1:100) in culture medium (Promega) was added to each well and the luminescence of the renilla luciferase was measured after 20 min incubation at RT. For data analysis, the Firefly luciferase signal was normalized to the renilla luciferase signal and statistical significance was determined through a t-test and plotted using Graph Pad Prism. All sequences for primers can be found in the Supplementary Table 1.

### CRISPRi and CRISPRa in dCas9-U251-MG

Plasmid encoding Lenti-dCas9-KRAB-blast for CRISPRi was a gift from Gary Hon (Addgene plasmid #89567; http://n2t.net/addgene:89567) and the plasmid for CRISPRa was acquired from Addgene (plasmid #96917; https://www.addgene.org/96917/). The plasmids were packaged into a lentivirus and U251-MG were transduced with 200 µL of each virus. Two days later, blasticidin (Gibco) at a concentration of 10 µg/mL was supplied to the culture medium and cells were continuously kept under antibiotic selection and the resulting clonal cells are denoted as dCas9-KRAB U251-MG for CRISPRi and dCas9-VPR U251-MG for CRISPRa. Plasmids containing guide RNAs (gRNAs) against each target were produced and lentivirally packaged. After 10 days of selection with blasticidin (Gibco) containing medium, 200.000 dCas9-KRAB U251-MG or dCas9-VPR U251-MG cells were seeded into a 6-well plate. One day later, cells were transduced with 15 µL of viruses containing the gRNAs against each target. As controls, two different NT constructs for CRISPRi and one for CRISPRa and a *PRNP* targeting construct were used. After 72 hours for CRISPRi and 24 hours for CRISPRa, cell media was replaced with fresh medium containing 1 µg/mL puromycin (Gibco) to select for transduced cells. Cell media was replenished once more with puromycin containing medium and selection of transduced cells was terminated after a total selection duration of 5 days for CRISPRi and 3 days for CRISPRa in culture. Subsequently, cells were washed once in PBS and lysed for downstream analysis by TR-FRET and immunoblotting as well as quantitative real-time polymerase chain reaction (qRT-PCR). For TR-FRET and immunoblotting, cells were lysed by scraping in a total volume of 80 µL of lysis buffer (containing 50 mM Tris-HCl pH 8, 150 mM NaCl, 0.5% sodium deoxycholate, 0.5% Triton-X 100, Sigma) and processed for immunoblotting as described in this manuscript under the PUM1 validation subchapter. A BCA assay was used for normalization to total protein concentration prior to FRET and Western Blot analyses. A BCA assay was used to assess the total protein concentration of the samples and the sample volume was adapted to achieve the same protein amount for subsequent FRET and Western Blot analyses. To quantitate PrP^C^ levels with a TR-FRET reaction the same antibodies were used as for the screening. In detail, 10 µl of lysates were manually transferred to a 384-well opaque OptiPlate (Perkin Elmer) in quadruplicate wells and supplied with APC-POM1 and EU-POM2 (5 and 2.5 nM final concentration, respectively diluted in LANCE buffer). Plates were incubated overnight at 4°C followed by centrifugation and TR-FRET measurements were done with the same parameters used for the screening workflow. qRT-PCR was followed as described in the previous PUM1 validation section of this manuscript following the same protocol. For CRISPRi, each sample was measured for three housekeeping genes (ACTB, GUSB and TBP) for normalization as well as their own respective primers to assess efficiency. For the final 9 hits, PRNP mRNA was assessed as well. For CRISPRa, each sample was measured for ACTB for normalization as well as their own respective primers to assess efficiency. qRT-PCR data was analyzed using the 2^-ΔΔCT^ method and visualized using GraphPad Prism. All primer sequences are listed under Supplementary Table 1.

### RNA-Sequencing in the U251-MG cell line

RNA extraction was performed using the RNeasy Mini Kit (Qiagen) according to the manufacturer’s instructions. The libraries were prepared following Illumina TruSeq stranded mRNA protocol. The quality of the RNA and final libraries was determined using an Agilent 4200 TapeStation System. The libraries were pooled equimolecularly and sequenced in an Illumina NovaSeq6000 sequencer (single-end 100 bp) with a depth of around 20 Million reads per sample. The experiment was run in triplicates. For thresholding for non-expressed or low expressed genes before CRISPRi validation, first the average normalized counts for triplicates were calculated and a cut-off value of 25 was used (Supplementary Table 1).

## Acknowledgements

AA is the recipient of an Advanced Grant of the European Research Council and grants from the Swiss National Research Foundation, the Nomis Foundation, the Swiss Personalized Health Network (SPHN, 2017DRI17), and a donation from the estate of Dr. Hans Salvisberg. DH is the recipient of a ZNZ Forschungskredit Candoc grant (FK-18-031). We would like to thank Dr. Emilio Yangüez and Dr. Maria Domenica Moccia and the Functional Genomics Center Zurich (FGCZ) for their help with the RNA-Sequencing experiment. We thank Irina Abakumova and Rita Moos for their technical help.

## Author Contributions

Conceived and designed the experiments: DH, MA, AA. Supervised the study: SH, PH, AA. Contributions to experimental work; siRNA primary screen: DH, MA, DP, siRNA reformatting: DH, MA, DP, ME, siRNA secondary screen and CRISPRi/a experiments: DH, MA, bioinformatics: DH, MA, ES, AC, design and cloning of CRISPRi/a guides: JAY, AS, virus packaging, DH, MA, KM, experimental planning and development for smNPC-derived neurons: AD, smNPC-derived neuron experiments: AD, DH, MA, PUM1 experiments: DH, MA, BD, qPCR experiments: DH, MA, BD, ME. Wrote the paper: DH, MA, AA.

## Supplementary materials

**Supplementary Figure 1:**
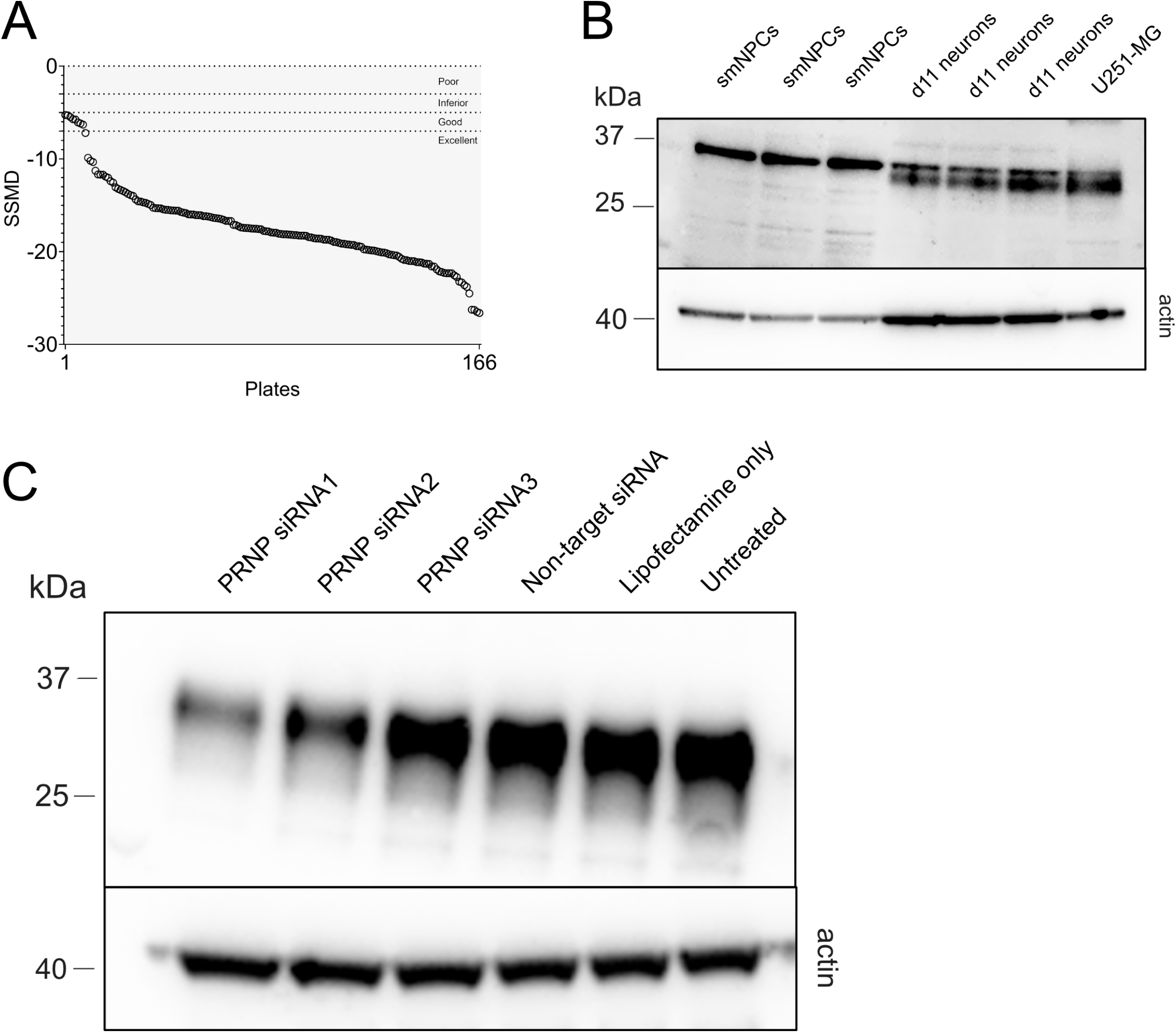
Additional quality control of primary screening and establishment of siRNA treatment for smNPC-derived neurons. **A)** Strictly standardized mean difference of each plate from the primary screening reporting the separation of the positive and negative controls. Range of the quality metrics (poor to excellent) is reported based on the highest level of stringency (Zhang, 2011). **B)** Endogenous PrP^C^ levels in human smNPCs and smNPC-derived neurons (day 11 post differentiation) assessed through immunoblotting. U251-MG was used as a positive control. The anti-PrP antibody POM2 was used for detection of PrP^C^. **C)** Western blot analysis of 11 days old smNPC-derived neurons upon transfection with three distinct *PRNP* targeting siRNAs and a non-targeting siRNA as well as Lipofectamine only and untreated control. *PRNP* siRNA 1 and 2 lead to a marked decrease of PrP^C^ levels 6 days after transfection.

**Supplementary Figure 2:**
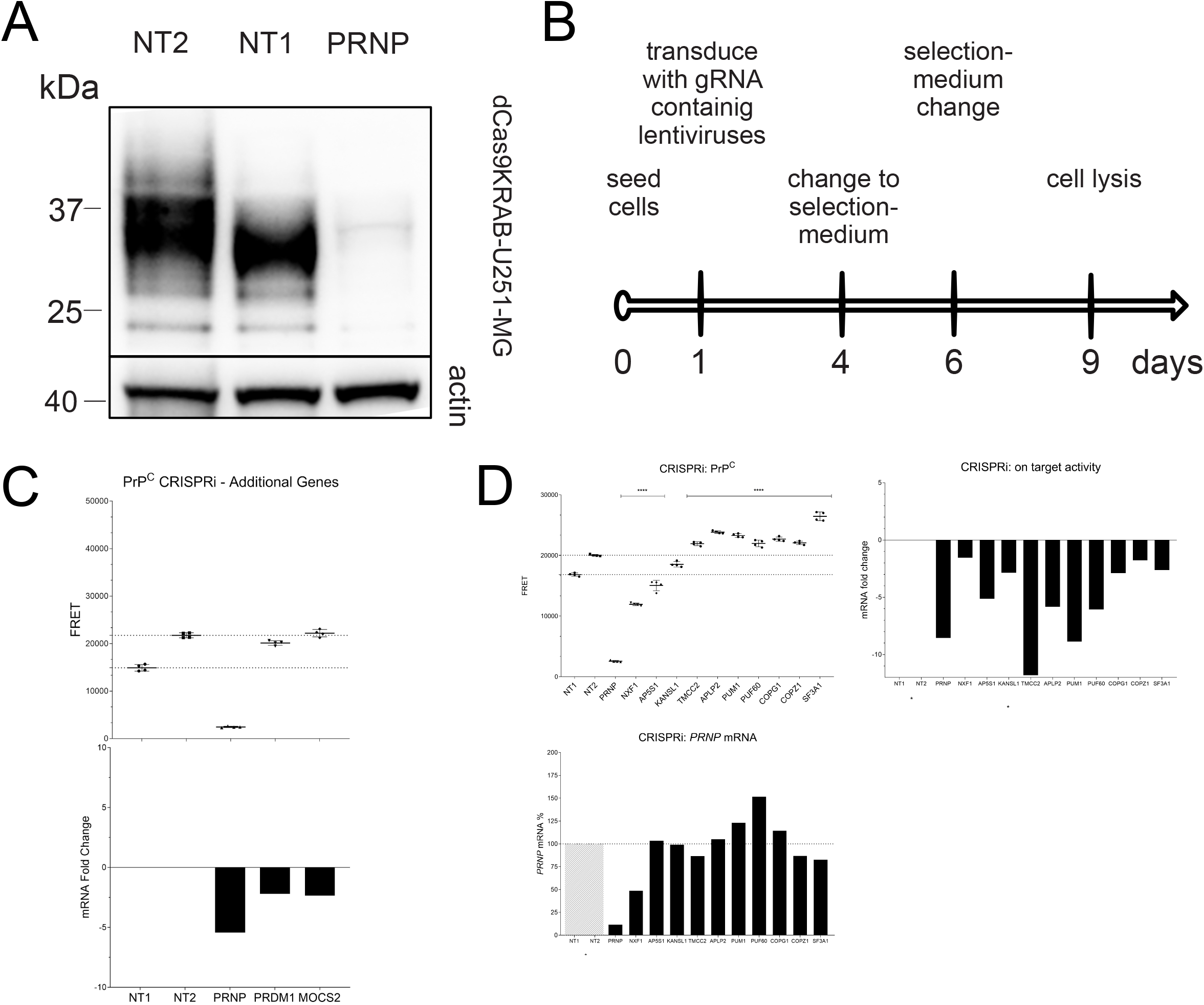
Establishment, experimental setup and results of CRISPRi in dCas9KRAB-U251MG cells. **A)** Efficacy of CRISPRi in dCas9-KRAB-U251-MG by lentivirally transduce a construct containing four single guides against *PRNP* as well as two non-targeting controls assessed through immunoblotting. After CRISPRi treatment followed by antibiotic selection for five days an evident reduction of PrP^C^ levels is apparent. **B)** Schematic depicting the setup of the CRISPRi experiments in dCas9-KRAB-U251-MG cells. **C)** Two candidates tested independently. PrP^C^ protein levels of dCas9KRAB-U251-MG cells transduced with gRNA CRISPRi lentiviruses against both targets. Mean values ± SD (n=4 technical replicates) are shown. CRISPRi activity measured by the mRNA level of the target gene in comparison to a negative control sample (NT1) after normalization to the housekeeping gene ACTB. **D)** Panels depicting the on-target efficiency of CRISPRi downregulation on each target assessed by qRT-PCR five days post-selection, the effect of each target on *PRNP* mRNA levels in the same experiment and the effect of CRISPRi for all targets assayed on PrP^C^ levels.

**Supplementary Figure 3:**
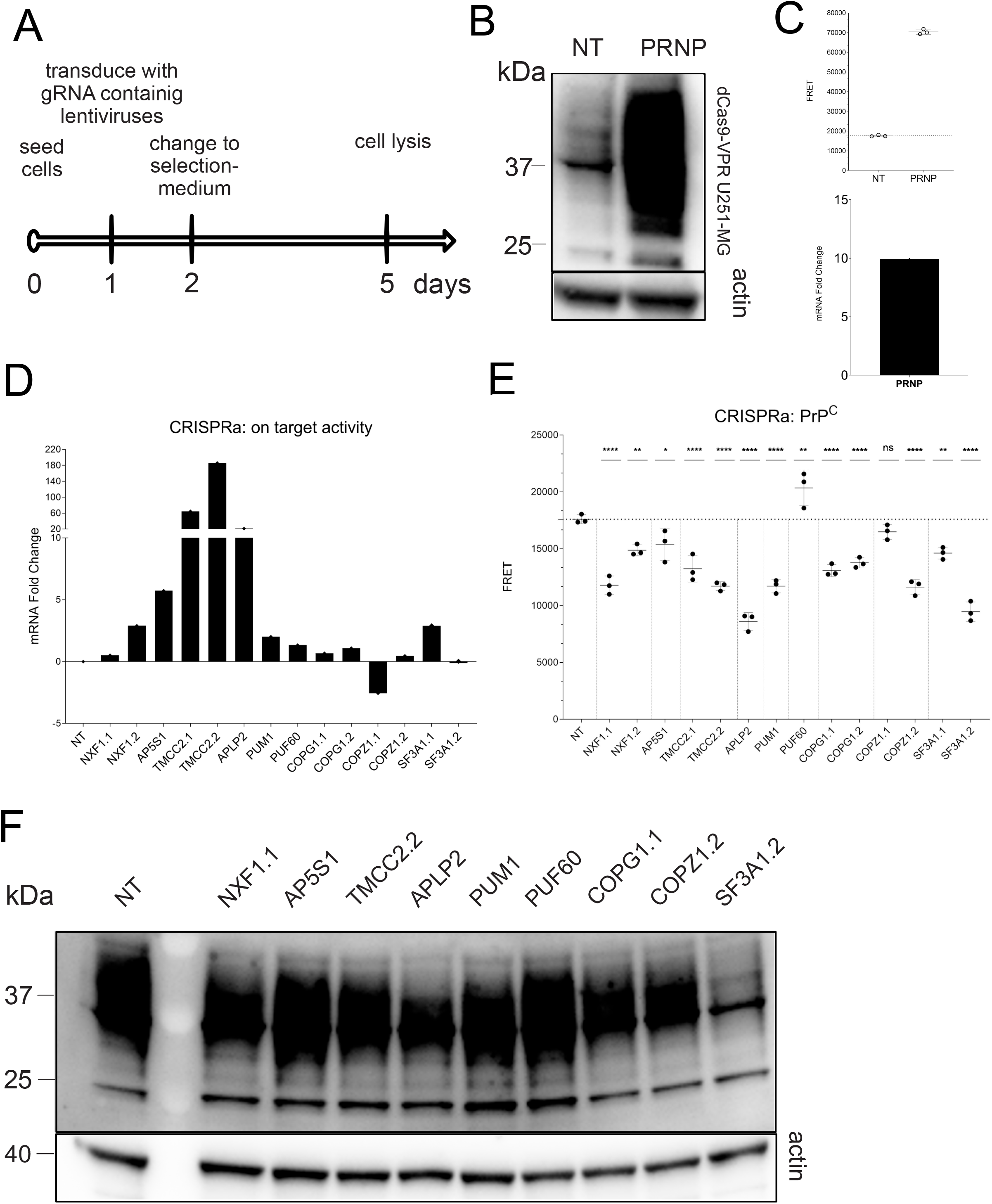
Establishment, experimental setup and results of CRISPRa in dCas9VPR-U251MG cells. **A)** Schematic depicting the setup of the CRISPRa experiments in dCas9-VPR-U251-MG cells. **B)** Efficacy of CRISPRa in dCas9-VPR-U251-MG by lentivirally transduce a construct containing gRNAs against *PRNP* as well as a non-targeting control assessed through immunoblotting. **C**) Additional quantification of B using TR-FRET. qRT-PCR of the same setup as in B Δ t values were normalized to those of *ACTB*. After CRISPRa treatment followed by antibiotic selection for three days an increase in PrP^C^ levels is apparent. **D)** CRISPRa efficiency for each gene measured by qRT-R after selection for three days Δ t values were normalized to those of *ACTB*. **E)** Effect of CRISPRa for all targets assayed on PrP^C^ levels measured by TR-FRET. **F)** Western blot analysis of representative samples from E. Antibody POM2 was used for detection of PrP^C^. * p ≥ 0.05, ** p ≥ 0 01, **** p ≥ 0 0001, n.s. = non-significant (Dunnett’s multiple comparisons test).

**Supplementary Table 1, Tab 1: Whole genome-RNAi screen results**. Summary excel sheet for the primary screen dataset involving SSMD, log2fold change and p-values for each replicate. Second tab summarizing the 743 candidates that were validated in the secondary screening round. Third tab summarizes the effects of the final nine hits.

**Supplementary Table 1, Tab 2: RNA-Seq normalized counts for U251-MG**.

**Supplementary Table 1, Tab 3: Sequences of primers used in the study**.

**Supplementary Table 1, Tab 4: MAGMA and VEGAS scores for the comparison of PrP**^**C**^ **hits and sCJD GWAS dataset**.

